# Bacterial Argonaute proteins aid cell division in the presence of topoisomerase inhibitors in *Escherichia coli*

**DOI:** 10.1101/2022.09.13.507849

**Authors:** Anna Olina, Aleksei Agapov, Denis Yudin, Anton Kuzmenko, Alexei A. Aravin, Andrey Kulbachinskiy

## Abstract

Prokaryotic Argonaute (pAgo) proteins are guide-dependent nucleases that function in host defense against invaders. Recently, it was shown that TtAgo from *Thermus thermophilus* also participates in the completion of DNA replication by decatenating chromosomal DNA. Here, we show that two pAgos from cyanobacteria *Synechococcus elongatus* (SeAgo) and *Limnothrix roseae* (LrAgo) act as DNA-guided DNA nucleases in *Escherichia coli* and aid cell division in the presence of the gyrase inhibitor ciprofloxacin. Both pAgos are preferentially loaded with small DNA guides derived from the sites of replication termination. The amount of pAgo-associated small DNAs (smDNAs) from the termination sites is increased in the presence ciprofloxacin, suggesting that smDNA biogenesis depends on DNA replication and is stimulated by gyrase inhibition. Ciprofloxacin also enhances asymmetry in the distribution of smDNAs around Chi-sites, indicating that it induces double-strand breaks that serve as a source of smDNA during their processing by RecBCD. While active in *E. coli*, SeAgo does not protect its native host *S. elongatus* from ciprofloxacin. These results suggest that pAgo nucleases help to complete replication of chromosomal DNA by targeting the sites of termination, and may switch their functional activities when expressed in different host species.

## INTRODUCTION

Argonaute (Ago) proteins are an evolutionary conserved family of programmable nucleases that are found in all three domains of life (1-3). Eukaryotic Argonautes (eAgos) participate in RNA interference and use small RNA guides to recognize RNA targets (4-7). This is followed by target RNA cleavage through an intrinsic endonucleolytic activity of eAgo or by recruitment of accessory factors, resulting in post-transcriptional or transcriptional gene silencing (8-11). When first identified in bacteria and archaea, prokaryotic Argonautes (pAgos) served as models to study the structure and biochemical properties of Ago proteins (3, 12-18), but their functional activities in host species remained unknown. Phylogenetic analysis demonstrated that pAgos are much more diverse than eAgos, and only a smaller part of them are catalytically active (1, 2, 19, 20). This analysis also revealed widespread horizontal transfer of pAgos among bacterial and archaeal species, suggesting that they can function in various genetic contexts and environments.

*In vitro* studies of several pAgo proteins demonstrated that their primary target is DNA rather than RNA. Recognition of DNA targets by pAgos can be guided by small DNAs, as observed for most studied pAgos, or by small RNAs (21-30). These findings were corroborated by *in vivo* analysis of several DNA-targeting pAgo proteins in bacterial cells, which revealed that pAgos preferentially recognize foreign DNA such as plasmids, mobile elements and phages. In particular, it was shown that RsAgo from *Rhodobacter sphaeroides*, TtAgo from *Thermus thermophilus*, PfAgo from *Pyrococcus furiosus*, and CbAgo from *Clostridium butyricum* decrease plasmid DNA content and transformation efficiency, and CbAgo counteracts phage infection (22, 26, 28, 29, 31). Accordingly, these pAgos are preferentially loaded with guide molecules corresponding to plasmid or phage sequences during their expression in *E. coli* (26, 29, 31). As demonstrated for CbAgo, generation of small guide DNAs from foreign genetic elements depends on both the catalytic activity of pAgo itself and the action of cellular nucleases (31). These studies suggested that catalytically active pAgos perform targeted degradation of foreign DNA, even if they are expressed in heterologous cells.

Recent studies suggested that elimination of invader DNA may not be the sole function of pAgos. In particular, TtAgo was shown to increase the resistance of its host bacterium *T. thermophilus* to ciprofloxacin, an inhibitor of DNA gyrase that impairs DNA replication and prevents normal cell division (32). TtAgo was shown to target the region of replication termination and participate in decatenation of chromosomal DNA, thus helping to complete DNA replication when the gyrase function is inhibited (32). However, it remained unknown whether this function in cell division is conserved among other DNA-targeting pAgos.

Here, we have analyzed two pAgo proteins from mesophilic cyanobacteria, SeAgo from *Synechococcus elongatus* and LrAgo from *Limnothrix roseae*. SeAgo is more closely related to TtAgo (35.7% identity in the MID and PIWI domains), while LrAgo is more distant from it on the phylogenetic tree (25.8% identity) (19). Both SeAgo and LrAgo are DNA-guided DNA nucleases that can perform precise cleavage of target DNA *in vitro* (24, 25). SeAgo was previously shown to interact with small guide DNAs in its native host *S. elongatus*, but without obvious target specificity (25). Here, we have found that, when expressed in a heterologous *E. coli* host, both SeAgo and LrAgo are loaded with small DNAs corresponding to the sites of replication termination. Furthermore, both SeAgo and LrAgo increase *E. coli* resistance to the gyrase inhibitor ciprofloxacin, suggesting that targeting of termination sites by pAgos may aid DNA replication in various prokaryotic species.

## RESULTS

### Cyanobacterial pAgos rescue *E. coli* growth in the presence of topoisomerase inhibitors

To explore whether SeAgo and LrAgo can affect DNA replication and cell division, we expressed them in the heterologous *E. coli* system. The SeAgo and LrAgo genes were cloned under the control of an arabinose-inducible promoter in pBAD-based vectors (Fig. 1A). Western blots confirmed that both proteins were expressed after the addition of arabinose (Fig. S1A). We then studied their effects on cell growth and resistance to gyrase inhibition, and further analyzed their DNA specificity *in vivo* (Fig. 1).

**Fig. 1.**
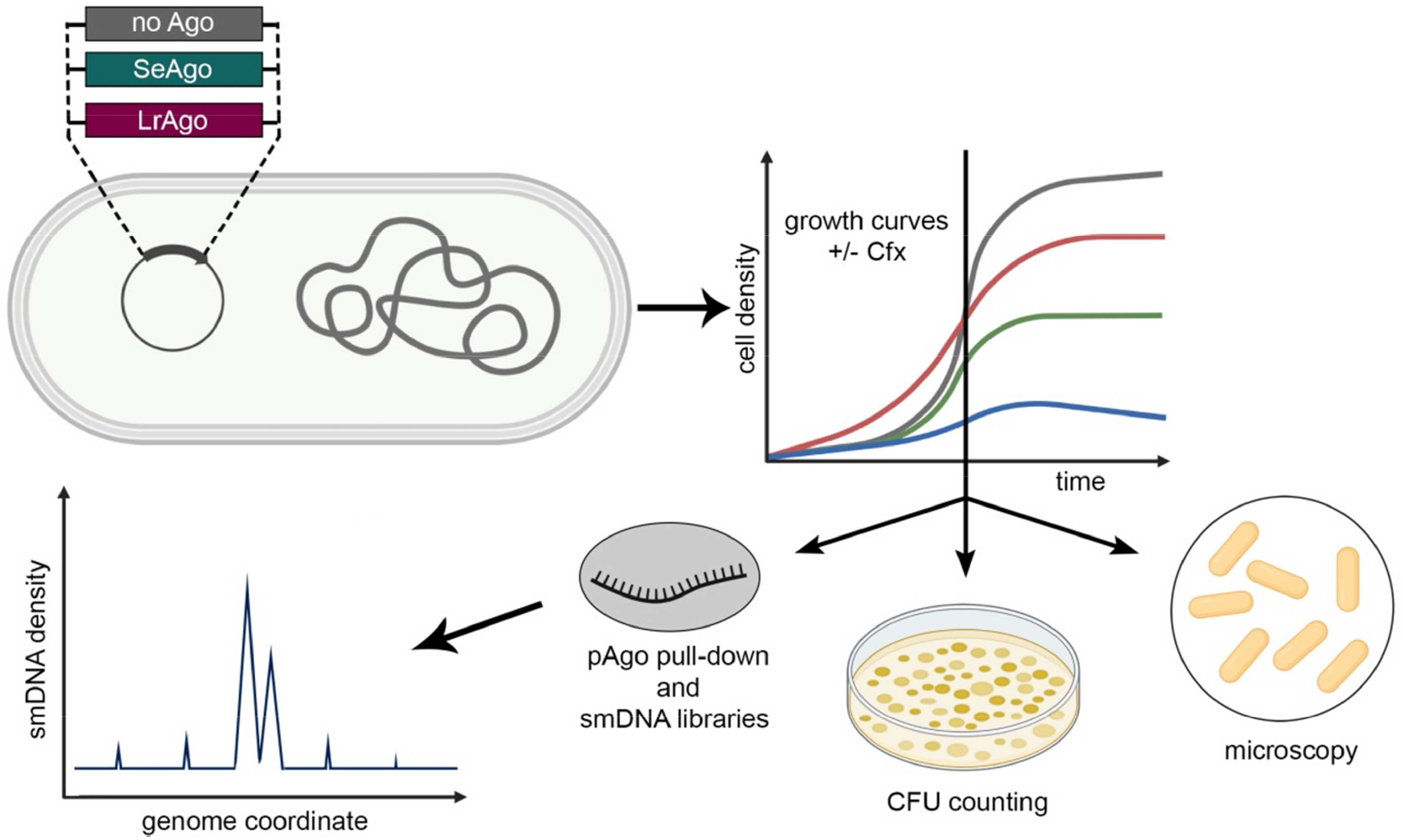
Analysis of pAgo functions in *E. coli.* *E. coli* strains expressing plasmid-encoded SeAgo or LrAgo or containing a control empty plasmid were grown in the absence or in the presence of ciprofloxacin (Fig. 2), followed by cell microscopy (Fig. 3), CFU counting (Fig. S2), and analysis of pAgo-associated smDNAs (Figs. 4-6, S3, S4).

We first determined the range of sublethal concentrations of ciprofloxacin in the absence of pAgo proteins, by measuring the kinetics of cell growth (OD_600_ curves) of an *E. coli* strain containing an empty expression plasmid. The cell growth was partially inhibited at 0.3-0.5 μg/ml of ciprofloxacin and was almost completely inhibited at 3 μg/ml of ciprofloxacin (Fig. S1B). We then tested the effects of the pAgo proteins on cell growth. LrAgo and SeAgo did not affect the growth kinetics in the absence of ciprofloxacin, however, both proteins protected the cells from the sublethal concentration of ciprofloxacin (0.5 μg/ml). LrAgo partially rescued cell growth, while SeAgo was able to restore cell growth completely (Fig. 2A and S3A). These experiments suggested that both pAgos can help the cells overcome the inhibitory effects of ciprofloxacin on DNA replication.

**Fig. 2.**
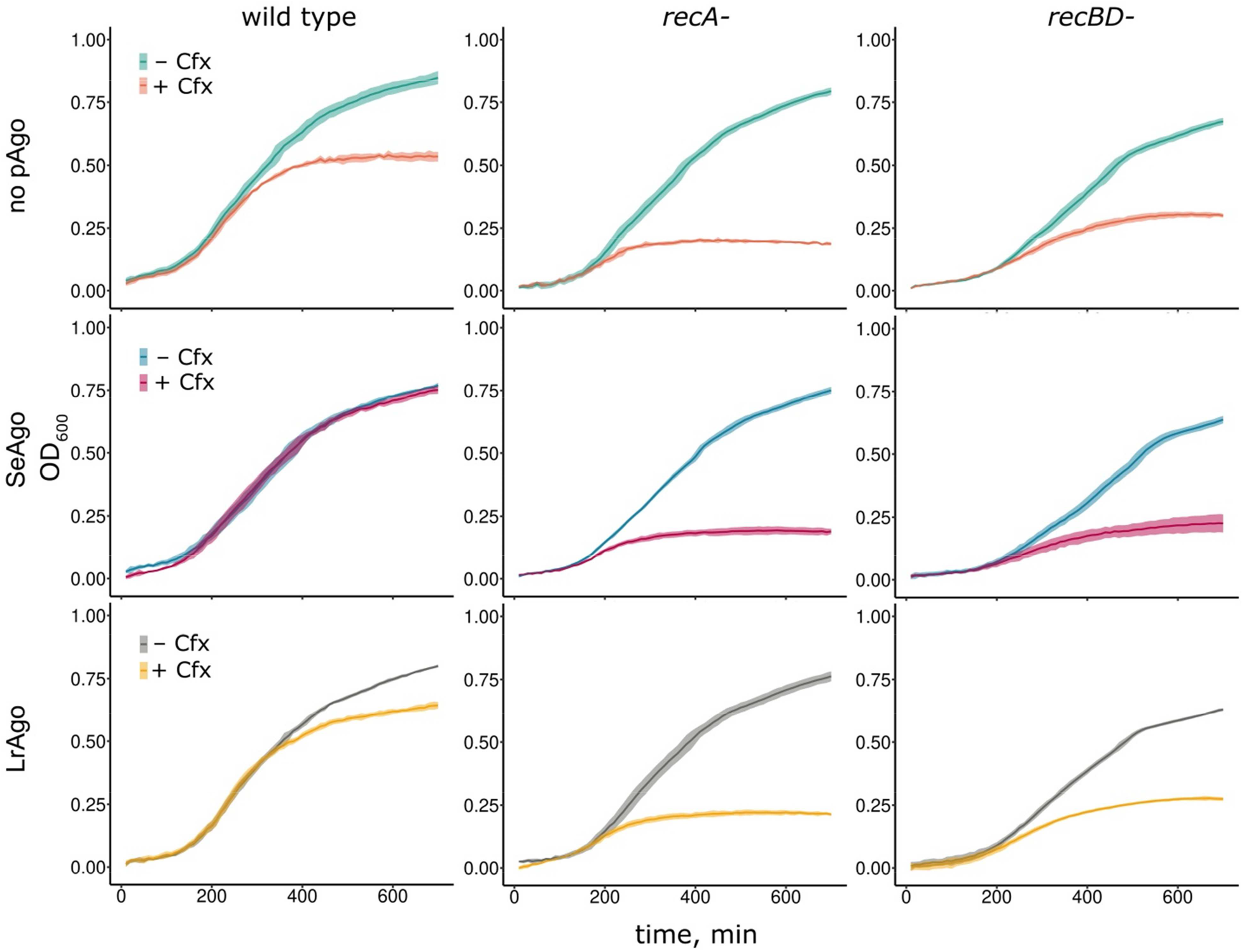
Growth of *E. coli* strains lacking or containing pAgos in the absence and in the presence of ciprofloxacin. The experiment was performed with wild-type (A) and *rec*-minus strains (B) in a plate reader. Ciprofloxacin was added to 0.5 μg/ml when indicated. Averages and standard deviations from 3 biological replicates are shown.

Previous experiments with TtAgo suggested that it helps to decatenate chromosomes by introducing double-strand breaks in the genomic DNA of *T. thermophilus* (32). If SeAgo and LrAgo acted by a similar mechanism, double-strand breaks generated during this process should subsequently be repaired by homologous recombination, depending on the RecA protein and the RecBCD helicase-nuclease involved in double-strand break processing (33-35). To test whether this was the case, we measured the effects of ciprofloxacin and pAgos in *E. coli* strains with deletions of *recA* and *recB/recD*. As expected, ciprofloxacin had stronger effects on the growth of these strains in comparison with wild-type cells (Fig. 2B). Notably, SeAgo and LrAgo did not stimulate the growth of *recA-*minus and *recBrecD*-minus strains in the presence of ciprofloxacin (Fig. 2B), suggesting that the function of the pAgo proteins depends on the homologous recombination machinery.

### pAgos suppress the effects of ciprofloxacin on cell division and morphology

The observed effects of ciprofloxacin on the density of cell cultures may not directly correspond to changes in the number of viable bacteria, because problems in cell division caused by the antibiotic induce formation of multinucleated cells (36) (see Discussion). We therefore directly compared the number of colony forming units (CFU) and analyzed cell morphology in bacterial cultures grown in the absence and in the presence of ciprofloxacin. After 4.5 hours of growth, when the effects of ciprofloxacin just became visible on the growth curves obtained by optical density measurements (Fig. 2), ciprofloxacin dramatically decreased CFU numbers in the wild-type *E. coli* strain (20- to 360-fold in three replicate experiments) (Fig. S2). Microscopy analysis revealed that the inhibition of cell division by ciprofloxacin caused disappearance of individual cells and formation of long multinucleated filaments (Fig. 3A, top panels).

**Fig. 3.**
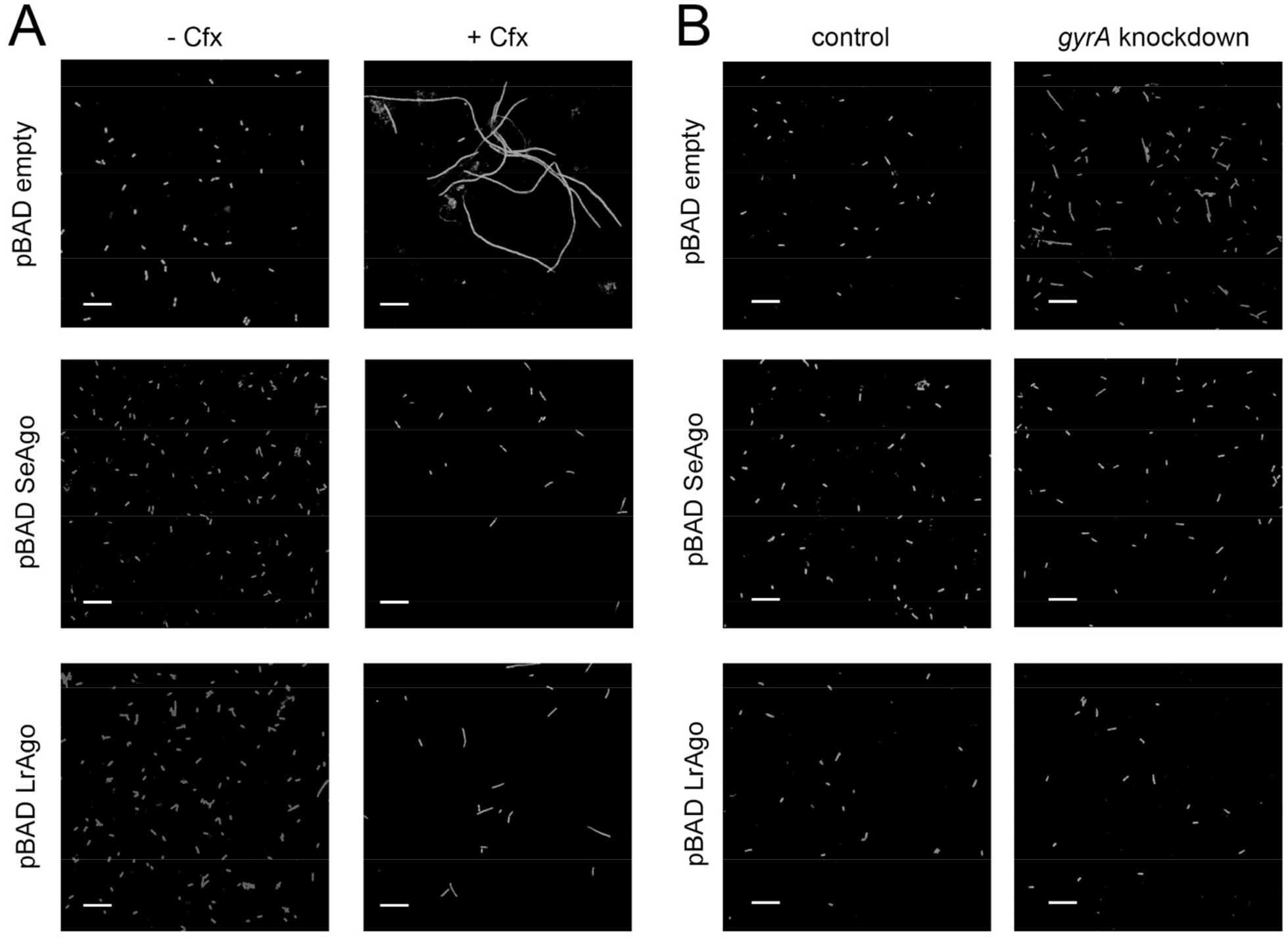
Effects of pAgo expression and gyrase inhibition on *E. coli* cell morphology. (A) *E. coli* cells lacking or containing pAgos were grown in the absence (left) or in the presence (right) of ciprofloxacin. The samples were taken at 4.5 hours from the cultures shown in Fig. 2. (B) Effects of gyrase (*gyrA*) knockdown on cell morphology. Fluorescence microscopy after acridine orange staining. The scale bar is 10 μm.

We then analyzed the effects of pAgo expression on the number of viable bacteria. In the absence of ciprofloxacin, CFU numbers were similar in control and pAgo-expressing strains (Fig. S2). In contrast, in the presence of ciprofloxacin CFU numbers were strongly increased in the strains expressing pAgos. This effect was especially prominent in the case of SeAgo; for this strain, ciprofloxacin decreased cell numbers only 3-16-fold in comparison with up to 360-fold inhibition observed in the control strain (Fig. S2). Microscopy analysis showed that expression of both SeAgo and LrAgo resulted in disappearance of long filaments induced by ciprofloxacin and increased the number of individual cells (or short filaments in the case of LrAgo) (Fig. 3A, middle and bottom panels).

Since ciprofloxacin primarily targets gyrase, we analyzed the effects of gyrase knockdown on cell morphology in control cells and upon pAgos expression. To silence gyrase expression, catalytically-inactive dCas9 and sgRNA corresponding to the beginning of the coding region of the *gyrA* gene were expressed in the *E. coli* strains lacking or containing pAgos. Control experiments demonstrated that only low level of gyrase knockdown could be achieved by this approach (10-20% decrease in the mRNA levels measured by quantitative PCR). However, this was sufficient to observe formation of short cell filaments by microscopy (Fig. 3B, top panels). Similarly to the experiments with ciprofloxacin, these filaments disappeared in the presence of pAgos (Fig. 3B, middle and bottom panels). Together, these results suggest that pAgos can aid cell division in *E. coli* cells when gyrase is inhibited by either ciprofloxacin or transcriptional knockdown, indicating that they may help to complete DNA replication impaired by topoisomerase deficiency.

### pAgo-bound smDNAs are generated in RecBCD-dependent manner and are enriched in the termination region of the chromosome

Previous analysis of small guide DNAs (smDNAs) associated with TtAgo in its native host demonstrated that they are highly enriched around the region of replication termination. This led to the suggestion that TtAgo may facilitate decatenation of chromosomal DNA by targeting the *ter-*region of the chromosome and possibly introducing double-strand breaks in this region (32). To explore whether SeAgo and LrAgo have preference for specific genomic regions or sequence motifs, we checked whether they were loaded with smDNAs during their expression in the heterologous *E. coli* host. Electrophoretic analysis of nucleic acids co-purified with SeAgo and LrAgo revealed that both pAgos were associated with 14-18 nt smDNAs (Fig. S3B). We sequenced libraries of pAgo-associated smDNAs obtained from late logarithmic or stationary bacterial cultures (for 5.5 and 12.5 h time points, Fig. S3A) in the absence or in the presence of ciprofloxacin and analyzed the distribution of smDNAs along the chromosomal and plasmid DNA.

Sequence analysis of smDNAs confirmed that the majority of smDNA associated with SeAgo and LrAgo have the length of 15-19 nt and 14-19 nt, respectively (Fig. 4A, top). Except of slight preference for G at the first guide position in the case of SeAgo, no strong nucleotide biases were found along the guide length. The mean GC-content of smDNAs corresponded to genomic DNA of *E. coli* (∼51%), and it was only slightly increased at the first guide position for SeAgo and slightly decreased upstream of the guide 5’-end and around 10-15 guide nucleotides for LrAgo (Fig. 4A, bottom). Overall, this analysis suggests that both SeAgo and LrAgo have no specific motif preferences and can likely interact with guide DNAs of any sequence.

**Fig. 4.**
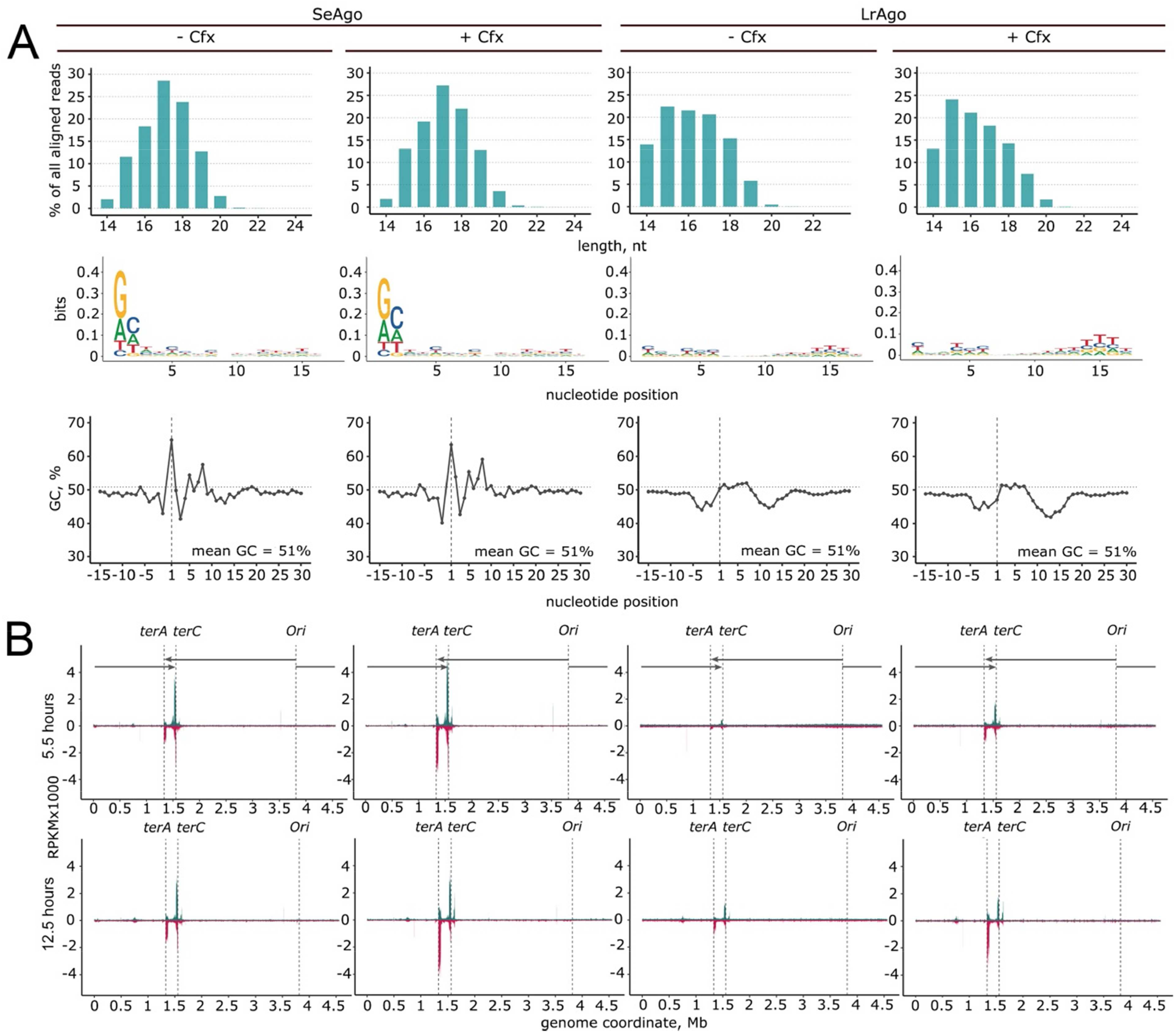
Whole-genome analysis of smDNAs associated with pAgos in *E. coli*. (A) Characteristics of pAgo-associated smDNAs obtained in the absence or in the presence of ciprofloxacin (for samples isolated at 5.5 hours of growth). (Top) Distribution of smDNA lengths for each smDNA library. (Middle) Nucleotide logos for different smDNA positions starting from the 5’-end. (Bottom) GC-content of the smDNA sequences and of the surrounding genomic regions. (B) Genomic distribution of smDNAs isolated from SeAgo and LrAgo in the absence or in the presence of ciprofloxacin at the logarithmic or stationary phases of growth (5.5 and 12.5 hours; see Fig. S3A for the growth curves). The numbers of smDNAs along the genomic coordinate are shown in reads per kilobase per million reads in the smDNA library (RPKM), individually for the plus (green) and minus (red) genomic strands. The *ori, ter* sites and the direction of replication are indicated.

Similarly to several previously studied pAgos, both SeAgo and LrAgo were enriched with smDNAs derived from plasmid DNA (11-14-fold enrichment over chromosomal DNA for SeAgo and 3-9-fold preference for LrAgo, after accounting for the relative replicon lengths and the plasmid copy number, 12 for pBAD). A similar bias for plasmid DNA was observed in the presence of ciprofloxacin (Table S1). Nevertheless, the majority of smDNA guides (78-95% in various experiments, Table S1) were derived from the *E. coli* chromosome indicating that genomic DNA is efficiently targeted by both pAgos.

The distribution of smDNA guides along the chromosome was highly uneven, with two large peaks at the *terA* and *terC* sites of replication termination for both pAgos (Fig. 4B, Fig. 5A). In addition, two smaller peaks were present at the next pair of *ter* sites, *terB* and *terD* (Fig. 5A). For SeAgo, similar targeting of the *ter* sites was observed at different stages of cell growth. For LrAgo, targeting of the *ter* sites was less efficient at the logarithmic stage but was increased in the stationary culture (Fig. 5A).

**Fig. 5.**
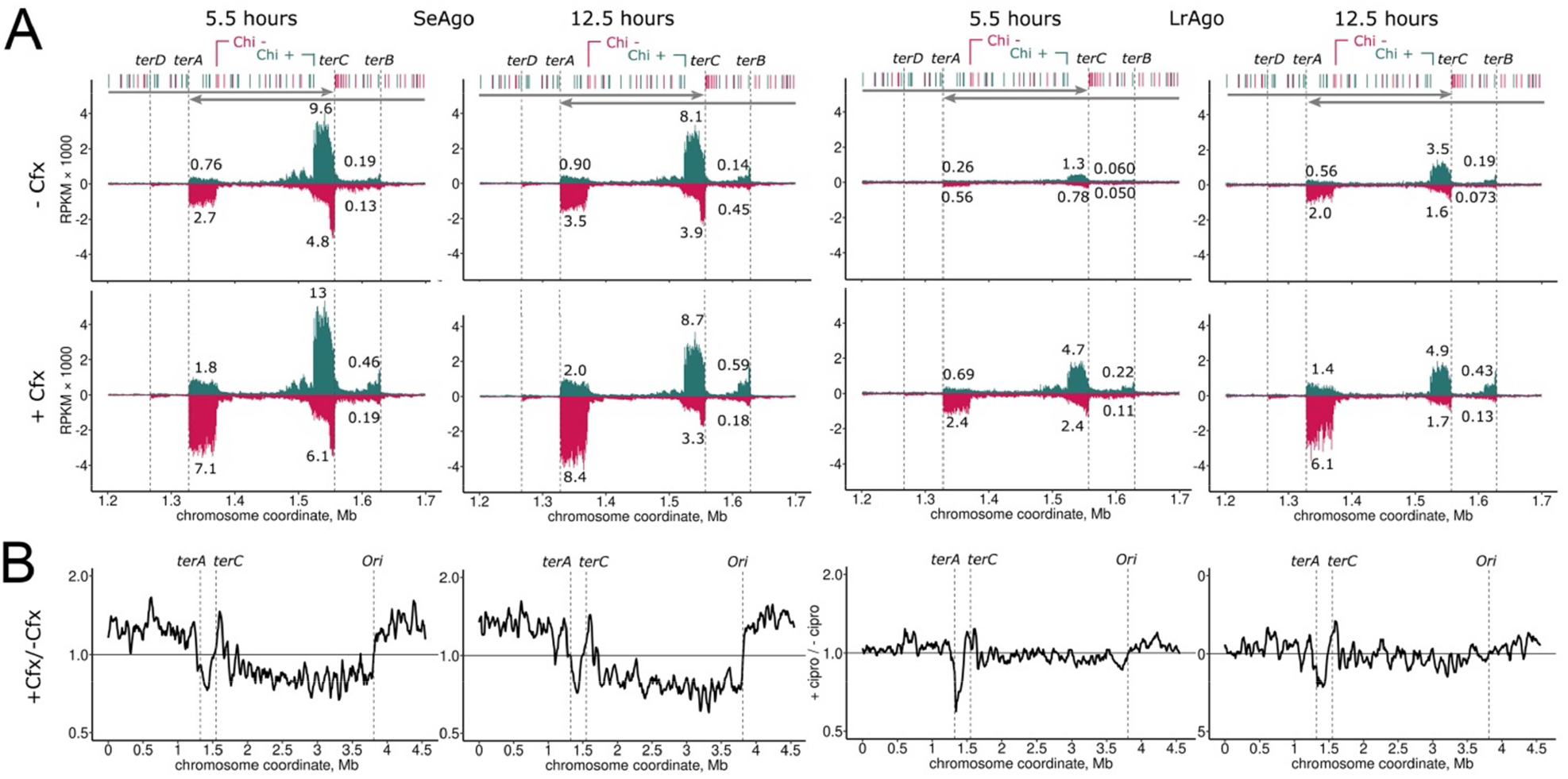
Targeting of the *ter* region by pAgos. (A) SmDNA densities in each genomic strand (plus strand, green; minus strand, red) in the *ter* region in *E. coli* cultures grown in the absence (top) and in the presence (bottom) of ciprofloxacin Positions of the *terA, terC, terD* and *terB* sites and the directions of replichores are shown. Chi sites in the plus (green) and minus (red) strands are shown above the plots. The closest Chi sites oriented toward *terA* (“Chi minus”), and *terC* (“Chi plus”) are indicated. SmDNA numbers are shown in RPKM in 1 kb windows. The numbers of smDNAs from each *ter* site, calculated for genomic regions between the *ter* site and the closest Chi site in the proper orientation, are shown as a percentage of the total number of smDNAs mapped to the whole genomic sequence in each smDNA library. (B) Changes in smDNA distribution across the chromosome caused by ciprofloxacin. pAgo-associated smDNAs were obtained from strains grown either in the presence or in the absence of ciprofloxacin. For each smDNA library, the ratio of smDNAs corresponding to the plus and minus genomic strands was first calculated, and the profiles obtained in the presence and in the absence of ciprofloxacin were then divided by each other. The resulting ratio shows changes in smDNA loading from the two genomic strands induced by ciprofloxacin.

The outer edges of the smDNA peaks precisely correspond to the *ter* motifs in chromosomal DNA bound by Tus protein (Fig. 5A). The inner borders of the peaks coincide with the properly oriented Chi-sites in each genomic strand (the closest sites in the plus strand for *terC* and in the minus strand for *terA*). Chi-sites (5′-GCTGGTGG-3′ in *E. coli*) serve as stop-signals for the RecBCD helicase-nuclease during processing of double-strand DNA breaks. This suggests that smDNAs are produced with participation of RecBCD from double-strand DNA ends that are formed after stalling of the replication fork at Tus-bound *ter* sites. In a further support for this notion, the ratio of smDNAs loaded from the plus and minus genomic strands in the *ter* region was asymmetric depending on the orientation of the *ter* sites. For both *terA* and *terC* (and *terB*), more smDNAs were produced from the DNA strand with the 3’-end oriented toward the *ter* site (from the minus genomic strand for *terA* and from the plus strand for *terC*; 2-3-fold more than from corresponding 5’-terminated DNA strands). This corresponds to the asymmetry in smDNA processing by RecBCD previously reported for another pAgo protein, CbAgo (31).

Targeting of the *ter* region by both SeAgo and LrAgo was increased in the presence of ciprofloxacin (Fig. 5A). Specifically, the fraction of smDNAs from *terA* was increased ∼2.6-fold and 3.9-fold for SeAgo and LrAgo, respectively, at the logarithmic stage of growth, and 2.4-fold and 2.9-fold at the stationary stage of growth (Fig. 5A). Furthermore, ciprofloxacin changed the relative sizes of the smDNA peaks at *terA* and *terC*. In the absence of ciprofloxacin, the peak at *terC* was larger than at *terA* for both pAgos at both stages of growth (Fig. 5A). This corresponds to the shorter length of the rightward replichore that terminates at *terC* and indicates that smDNAs at *ter* sites are likely produced during replication termination, which is more frequent at *terC* (31, 37). In contrast, the peaks at *terA* and *terC* were comparable in the presence of ciprofloxacin, because smDNA loading was stimulated to a higher extent at *terA* than at *terC* (Fig. 5A). For both pAgos, this effect became especially prominent in stationary cultures. This indicates that the relative frequencies of replication termination at *terA* and *terC* may be leveled in the presence of ciprofloxacin, or that the efficiency of smDNA processing in these regions may become less dependent on replication in these conditions.

To further explore the role of the RecBCD machinery in the processing of smDNAs guides, we calculated the distribution of smDNAs around Chi sites throughout the whole chromosome excluding the *ter* region. This metaplot analysis revealed that distribution pAgo-bound smDNA guides was asymmetric and dependent on the orientation relative to the Chi site. For the DNA strand co-oriented with Chi, the amounts of smDNAs derived from the 3’-side of Chi were much higher than from the 5’-side, with an abrupt drop immediately at the Chi sequence (Fig. 6, green). For the DNA strand that was oriented in the opposite direction relative to Chi, the changes in the amounts of smDNAs around Chi were much less pronounced (Fig. 6, gray). This indicates that smDNAs are preferentially generated from the 3’-terminated DNA strand during processing of double-strand ends by RecBCD, which stops at Chi sites.

**Fig 6.**
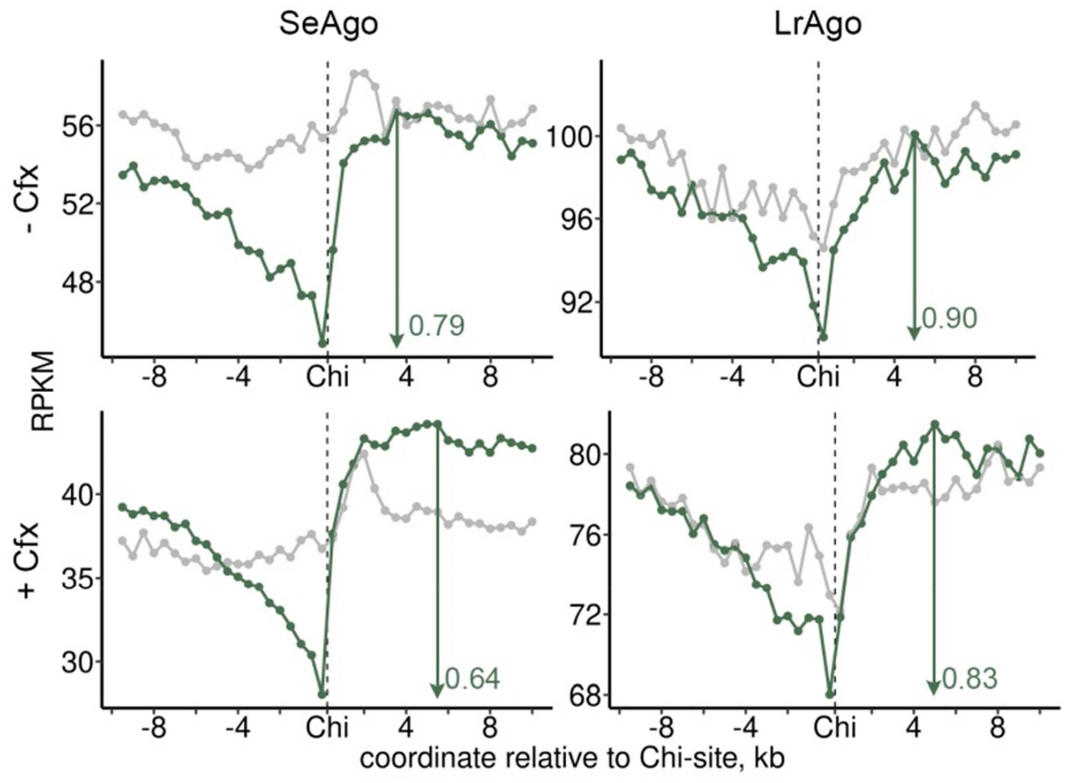
Asymmetry of smDNA distribution around Chi sites. Metaplots of the densities of smDNAs around Chi sites analyzed for smDNA libraries isolated from the logarithmic culture at 4.5 hours of growth (one of two replicate experiments). SmDNA numbers were independently calculated for the DNA strands co-oriented (green) and oppositely oriented (gray) relative to the Chi sequence (5’-GCTGGTGG-3’) for all Chi sites in both genomic strands except for the *ter* region (1.2-1.7 Mb chromosomal coordinates) and averaged in 0.5 kb windows.

For both pAgos, the asymmetry of smDNA loading around Chi sites was increased in the presence of ciprofloxacin. These changes were reproducible in two independent replicate experiments. In particular, the drop in the amounts of smDNAs upstream of the Chi motif was changed from 0.77 to 0.62 for SeAgo and from 0.89 to 0.81 for LrAgo in the logarithmic phase of growth (average from two replicates) (Fig. 6). This suggested that ciprofloxacin stimulates RecBCD-dependent processing of smDNAs, likely by increasing the number of DSBs formed in the chromosomal DNA due to inhibition of topoisomerases.

Finally, to explore possible connection of smDNA processing to replication, we compared the ratio of pAgo-bound smDNAs produced from the plus and minus genomic strands (Fig. 5B). In the case of SeAgo ciprofloxacin increased generation of smDNAs from the leading DNA strand in both replichores, resulting in an increased ratio (>1) of smDNAs corresponding to the plus and minus strands for the rightward replichore and a decreased ratio (<1) for the leftward replichore (Fig. 5B). In the *ter* region, this ratio was also changed in favor of the 3’-terminated strand at each *ter* site for both pAgos (<1 for *terA* and >1 for *terC*). This correlates with the observed increase in asymmetry of smDNA processing around Chi sites, which are mostly co-oriented with replication, and suggests that both changes result from increased stalling of replication forks in the presence of ciprofloxacin, followed by their processing by RecBCD.

### Distribution of pAgo-associated smDNAs relative to coding regions

*In vitro* experiments with several pAgo proteins, including TtAgo, CbAgo and LrAgo, demonstrated that their ability to process double-stranded DNA depends on DNA supercoiling (22, 24, 29). Our results indicate that gyrase inhibition, which changes the supercoiling state of the chromosome, also changes the distribution of smDNAs. Another factor that could potentially affect smDNA processing by changing DNA supercoiling is transcription, which induces positive supercoils in front of the moving RNA polymerase and negative supercoils behind of it (38-41). To explore the role of transcription in the targeting of chromosomal DNA by pAgos, we first compared the abundances of pAgo-associated smDNAs derived from each genomic strand for genes co-directed or oppositely directed relative to replication. No differences between these groups of genes were found for either SeAgo or LrAgo (Fig. S4A).

We then analyzed smDNA abundance in intergenic DNA regions for divergent, convergent and co-oriented gene pairs. Divergent and convergent gene orientation is associated with negative and positive DNA supercoiling in intergenic regions, respectively, which could affect smDNA production (38). No significant differences in the amounts of smDNAs were detected for the different types of gene pairs for both pAgos (Fig. S4B) suggesting that transcription is unlikely to have major effects on smDNA biogenesis.

### Analysis of the effects of ciprofloxacin and SeAgo on cell division in *S. elongatus*

The experiments presented above demonstrated that pAgo proteins from mesophilic cyanobacteria can suppress defects in DNA replication in *E. coli* caused by gyrase inhibition. To test whether these pAgos can have similar functions in their native species, we compared *S. elongatus* strains with the natural level of expression of SeAgo (wild-type strain), without SeAgo (ΔSeAgo), and with an increased level of SeAgo, expressed from a strong constitutive promoter (↑SeAgo). Titration of ciprofloxacin demonstrated that growth of all three strains in liquid culture was fully inhibited at ≥15 ng/ml of the antibiotic, indicating that *S. elongatus* is more sensitive to ciprofloxacin than *E. coli*. We then compared cell growth at sublethal ciprofloxacin concentrations. Wild-type and ΔSeAgo strains had identical growth kinetics in the absence and in the presence of 10 ng/ml of ciprofloxacin (Fig. 7A). In contrast, the growth of the strain with increased SeAgo expression was strongly inhibited in the presence of ciprofloxacin. Microscopy analysis revealed no changes in the cell number or morphology in either wild-type or ΔSeAgo strains in these conditions (Fig. 7B). Cell morphology also remained unchanged in the case of the SeAgo overexpressing strain, despite the lower number of cells observed in the presence of ciprofloxacin (Fig. 7B). Thus, ciprofloxacin inhibits *S. elongatus* growth without formation of multicellular filaments, suggesting that its mechanism of action in cyanobacterial cells may be different from *E. coli*. SeAgo does not increase the resistance of the wild-type strain of *S. elongatus* to ciprofloxacin in comparison with the deletion strain, while its overexpression is toxic in the presence of the antibiotic.

**Fig. 7.**
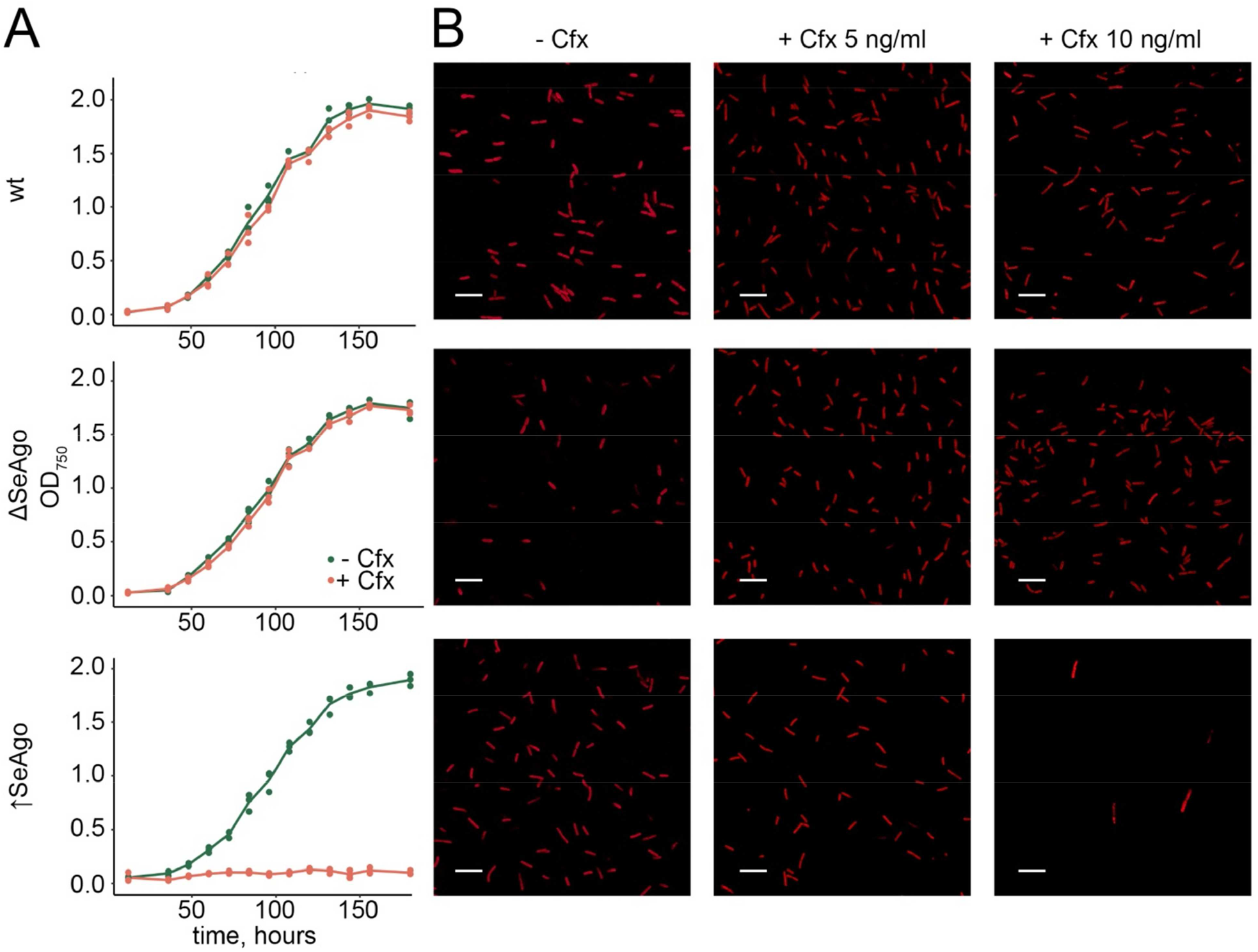
No effect of SeAgo on cell growth in *S. elongatus*. (A) Growth of *S. elongatus* strains with different levels of expression of SeAgo in the absence and in the presence of ciprofloxacin (10 ng/ml). Wt, wild-type strain; ΔSeAgo, strain without SeAgo; ↑SeAgo, strain with increased expression of SeAgo. Averages from 3 biological replicates are shown. (B) Analysis of cell morphology of the same *S. elongatus* strains in the absence or in the presence of ciprofloxacin (5 or 10 ng/ml). The samples were taken at 48 hours of growth and visualized by fluorescence microscopy.

## DISCUSSION

In contrast to eAgos that recognize RNA targets, the majority of studied pAgos preferentially target DNA *in vitro* and *in vivo*, suggesting that their mechanism of action is different from eAgos. Recently, a novel group of pAgos was shown to use DNA guides to recognize RNA targets (42, 43), but their functional activities *in vivo* remain to be investigated. Studied pAgos were shown to target foreign genetic elements in bacterial cells (22, 26, 28, 29, 31), suggesting that the defensive function of Ago proteins is conserved between prokaryotes and eukaryotes (44). At the same time, it was suggested that pAgos might play other roles, including the regulation of gene expression, participation in cellular suicide systems and DNA repair (20, 26, 45-48). Indeed, TtAgo from *T. thermophilus* was recently demonstrated to participate in separation of chromosomal DNA during replication (32). This activity of TtAgo became crucial for cell division when DNA gyrase - the sole type II topoisomerase in *T. thermophilus* - was inhibited by ciprofloxacin. It was shown that TtAgo is associated with small guide DNAs corresponding to the termination region of replication and can be co-precipitated with several proteins involved in DNA processing, including gyrase and factors involved in DNA recombination (AddAB, a RecBCD homolog in *T. thermophilus*). It was therefore proposed that TtAgo helps to decatenate daughter chromosomes by direct DNA cleavage and/or by recruiting accessory factors to the termination region (32).

Here, we have demonstrated that two pAgo proteins from mesophilic cyanobacteria, SeAgo and LrAgo, facilitate cell division and prevent formation of multinucleated cell filaments in the presence of ciprofloxacin in *E. coli*. The primary target of ciprofloxacin in *E. coli* is gyrase (49-52), and both pAgos also suppress a milder phenotype caused by dCas9 knockdown of gyrase expression in *E. coli*. Inhibition of gyrase is known to strongly affect replication by changing DNA supercoiling and integrity, and by introducing direct roadblocks to the moving replisomes (52-55). Thus, formation of cell filaments in the presence of ciprofloxacin can be explained by failure to separate daughter chromosomes due to their incomplete replication and decatenation, and by the induction of the SOS response after DNA damage, preventing cell division (55-60).

SeAgo and LrAgo may help the cells to complete DNA replication by direct targeting of chromosomal DNA through their nuclease activity. In support of this proposal, we found that both SeAgo and LrAgo preferentially target the replication termination region of the *E. coli* chromosome and are loaded with small guide DNAs corresponding to the *ter* sites. The stronger effects of SeAgo on cell growth and morphology in the presence of ciprofloxacin (Fig. 2 and Fig. 3) correlate with its higher loading with smDNAs from the *ter* sites and a stronger asymmetry of smDNA processing around Chi sites and across the chromosome. Recent work by Jolly et al. suggested that TtAgo may recruit gyrase and double-strand break repair factors to the *ter* region in *T. thermophilus* (32). In comparison, SeAgo and LrAgo are not expected to specifically interact with host-specific factors in heterologous *E. coli* suggesting that they directly participate in chromosomal DNA processing. In particular, pAgos may introduce single-strand or double-strand breaks in the *ter* region when loaded with corresponding smDNAs, thus decreasing the level of unfavorable positive DNA supercoiling between the converging replisomes and/or assisting decatenation of the sister chromosomes in the presence of topoisomerase inhibitors. DNA decatenation requires introducing double-strand breaks into chromosomal DNA, and the anti-inhibitory function of pAgos indeed depends on the homologous recombination machinery and is not observed in *E. coli* strains lacking the key components of double-strand break repair, RecBCD or RecA. Further experiments are required to decipher the interplay between pAgos and the recombination machinery in various species.

Similar enrichment of pAgo-associated smDNAs at the *ter* sites in the chromosomal termination region was previously observed for TtAgo in *T. thermophilus* (32) and for CbAgo expressed in *E. coli* (31). Thus, different pAgo proteins have similar chromosomal preferences, indicating that the ability to recognize the sites of replication termination may be a common feature of pAgo proteins from various branches of the pAgo tree. This specificity is likely determined by the mechanism of replication termination and may be different in various species. Indeed, a different pattern of smDNA distribution is observed for SeAgo in its native host *S. elongatus*, which lacks recognizable *ter* sites in the chromosome (25).

In the *ter* region, smDNAs loaded into pAgos are preferentially generated from 3’-ends of DNA strands oriented toward *ter* sites, and smDNA processing is confined to the areas between *ter* sites and the closest Chi sites. Furthermore, smDNA processing throughout the genome outside of the *ter* region also depends on the position and orientation of Chi sites. At the same time, smDNA biogenesis does not significantly depend on the orientation of transcription units relative to each other or to the direction of replication. The observed pattern of smDNA processing from the *ter* region and its dependence on Chi sites indicate that smDNAs are produced during asymmetric cleavage of the two DNA strands by RecBCD or other cellular nucleases that cooperate with RecBCD during DNA unwinding (31, 33-35). The efficiency of smDNA processing from the *ter* region relative to other regions of the chromosome and the asymmetry of smDNA processing around Chi sites are increased in the presence of ciprofloxacin, possibly as a result of increased levels of gyrase-mediated DNA fragmentation induced by the antibiotic. This may result in increased targeting of the *ter* region by guide-loaded pAgos and help to overcome problems with chromosomal DNA processing during final steps of replication.

Strikingly, while SeAgo protects *E*.*coli* cells from the toxic effects of ciprofloxacin, it does not defend its host strain of *S. elongatus*. Moreover, overexpression of SeAgo makes *S. elongatus* more susceptible to this antibiotic. This may be explained by a different pattern of chromosomal targeting by SeAgo in *S. elongatus* (25) in comparison with *E. coli* and/or by differences in the mechanism of inhibition of cell division by ciprofloxacin in cyanobacteria. In particular, cyanobacteria do not have classical SOS response characterized in *E. coli*, and *S. elongatus* apparently lacks its master regulator, the LexA repressor (61, 62). SeAgo may therefore increase DNA damage rather than aid DNA processing in *S. elongatus* cells treated with ciprofloxacin.

Natural functions of SeAgo in cyanobacterial cells remain to be investigated. We have previously shown that its deletion or overexpression do not affect the kinetics of cell growth under laboratory conditions (25). At the same time, loss of function of SeAgo was shown to increase the efficiency of plasmid DNA transfer (63, 64). SeAgo may therefore modulate natural transformation in its host bacterium. Our findings suggest that the functions of pAgo proteins may change upon their transfer between prokaryotic species and switch between cell protection, regulation of horizontal gene transfer, and processing and repair of chromosomal DNA.

## MATERIALS AND METHODS

### Plasmids and strains

All *E. coli* strains were isogenic to *E. coli* BL21(DE3). *recA-*minus and *recBD-*minus strains were obtained previously (31). *E. coli* cells were routinely cultivated in LB Miller broth (2% tryptone, 0.5% yeast extract, 1% NaCl, pH 7.0) with the addition of ampicillin (100 μg/ml) if needed. The gene of SeAgo (WP_011378069.1) was codon-optimized using IDT Codon Optimization Tool for expression in *E. coli*, synthesized by the IDT core facility, and cloned into pBAD-HisB in frame with the N-terminal His6-tag. The pBAD plasmid encoding LrAgo was obtained previously (24). *E. coli* strains were transformed with pBAD plasmids encoding each of the two pAgos or a control plasmid without pAgo, grown overnight in LB with ampicillin, diluted twice with 50% glycerol, aliquoted, frozen in liquid nitrogen and stored at −80°C.

### Ciprofloxacin treatment and analysis of growth kinetics in *E. coli*

To determine sublethal ciprofloxacin concentrations, overnight bacterial culture of BL21(DE3) carrying the control pBAD vector without pAgo genes was obtained from the frozen aliquoted culture and inoculated into 100 μl of fresh LB medium supplemented with ciprofloxacin (0, 0.1, 0.3, 0.5 μg/ml), 0.01% L-arabinose and ampicillin in 96-well plates. The plates were incubated at 300 rpm at 30 °C in a CLARIOSTAR microplate reader and cell density was monitored by measuring OD_600_ every 10 min. Three independent biological replicates were performed.

For analysis of the effects of pAgos on the growth kinetics, overnight bacterial cultures were obtained from frozen aliquoted cultures and inoculated into 100 μl of fresh LB medium supplemented with 0 or 0.5 μg/ml ciprofloxacin, 0.01% L-arabinose to induce pAgo expression and ampicillin in 96-well plates (TPP, flat bottom). The plates were incubated at 300 rpm at 30°C in a CLARIOSTAR microplate reader and cell density was monitored by measuring OD_600_ every 10 min. Three independent biological replicates were performed. Additionally, after 4.5 hours cells were harvested for Western blot analysis, microscopy and determination of CFU numbers. To calculate CFU, serial dilutions of the culture were plated on selective agar medium containing ampicillin.

### Western blotting

The levels of SeAgo and LrAgo expression were determined by Western blotting. *E. coli* cells expressing pAgos were harvested by centrifugation, the pellet was mixed with 1^×^ Laemmli sample buffer (120 mM Tris-HCl, 4% SDS, 4% β-mercaptoethanol, 10% glycerol, pH 6.8) and heated at 95 °C for 5 min, and the samples were resolved by electrophoresis in a 4–20% Tris-glycine gel (BioRad). Proteins were transferred onto a nitrocellulose membrane in Towbin transfer buffer (25 mM Tris, 192 mM glycine, 20% methanol) using semi-dry procedure at 25 V, 1 A for 30 min (BioRad Trans-Blot Turbo). The transfer membrane was washed in PBS (10 mM phosphate buffer, 137 mM NaCl, 2.7 mM KCl) for 5 min. The membrane was blocked with blocking buffer (PBS, Tween-20 0.1% (v/v), non-fat milk 5% (w/v) for 30 min at room temperature, and then incubated with anti-His6 monoclonal antibodies (1:1,000, Sigma) for 1 h at room temperature. The membrane was washed four times with PBST buffer (PBS, Tween-20 0.1% (v/v)), and incubated with HRP-conjugated anti-mouse secondary antibodies (1:10,000, Sigma) for 1 h at room temperature and washed again as described above. Antigen–antibody complexes were detected with Immobilon ECL Ultra Western HRP substrate (Millipore) on a Chemidoc XRS+ imager (BioRad).

### Preparative growth of *E. coli* and purification and sequencing of pAgo-associated smDNAs

Overnight bacterial cultures were obtained from frozen aliquoted cultures and inoculated into 500 ml of fresh LB medium supplemented with 0 or 0.3 μg/ml ciprofloxacin, 0.01% L-arabinose and ampicillin in 2 liter flasks. The flasks were incubated at 190 rpm at 30°C in an orbital shaker and cell density was monitored by measuring OD_600_ every 30 min. Two independent biological replicates were performed. The cells were harvested by centrifugation at 7,000g, 4 °C for 15 min after 5.5 and 12.5 hours of growth for protein pull-down. The cells were disrupted with a high-pressure homogenizer (EmulsiFlex-C5, Avestin) at 18000 psi. pAgos were pulled down using Co^2+^-Talon Metal Affinity Resin (Takara) as described previously (26). Eluted proteins were treated with Proteinase K for 30 minutes at 37°C, small nucleic acids were extracted with phenol-chloroform, ethanol-precipitated, dissolved in water and analyzed by PAGE as described (26).

Libraries for high-throughput sequencing of smDNAs were prepared according to the previously published splinted ligation protocol (26). Briefly, nucleic acids extracted from pAgos were treated with RNase A (Thermo Fisher), purified by PAGE, small DNAs (14-20 nt) were eluted from the gel in 0.4 M NaCl overnight at 21 °C, ethanol precipitated, dissolved in water, phosphorylated with polynucleotide kinase (New England Biolabs), and ligated with adaptor oligonucleotides using bridge oligonucleotides as described in (26). The ligated DNA fragments were purified by denaturing PAGE, amplified, and indexed by the standard protocol for small RNA sequencing (New England Biolabs). Small DNA libraries were sequenced using the HiSeq2500 platform (Illumina) in the rapid run mode (50-nucleotide single-end reads). The list of all sequenced smDNA libraries is presented in Table S1.

### Analysis of chromosomal distribution of smDNAs

Analysis of smDNA sequences was performed as described previously (31). After trimming the adaptors, reads shorter than 14 nt were removed with CutAdapt (v. 2.8). Bowtie (v. 1.2.3) was used to align the reads to the reference genomic DNA (Refseq accession number NC_012971.2) with no mismatches allowed. Genome coverage was calculated using BEDTools (v. 2.27.1) and custom Python scripts. Whole-genome coverage of each DNA strand was calculated in 1000 nt windows and normalized by the total number of mapped reads in the library, expressed as RPKM (reads per kilobase per million mapped reads). To calculate the percent of reads mapped to the regions around *ter*-sites, the number of smDNAs for each DNA strand from each region (from 1328000 to 1350000 for *terA*, from 1524000 to 1557000 for *terC*, and from 1626000 to 1629000 for *terB*) was divided by the total number of reads mapped to both strand of the genome. The ratio between the coverage of plus-strand and minus-strand from the ciprofloxacin-treated cell culture was divided by the same ratio obtained for the control library and plotted as a rolling mean (50 kb window, 10 kb step). To calculate smDNA distribution around Chi-sites, in genes and in intergenic intervals, the region of replication termination (1.2-1.7 Mb chromosomal coordinates) was removed from the analysis. Coverage of the regions around Chi-sites was calculated in 500-nt bins and then averaged. Gene borders were extracted from the Refseq annotation file. Intergenic intervals with length < 500 nt were classified into four groups depending on the relative orientation of the surrounding gene, and 100 nucleotides were additionally included from each side to each interval. SmDNA coverage of genes and intergenic intervals was calculated in RPKM. Nucleotide logos were calculated using reads longer than 16 nt, and reads longer than 17 nt were truncated to 17 nt from the 3’-end. All plots were generated in R (v. 3.6.3) using ggplot2 (v. 3.3.3) and ggseqlogo (v. 0.1) (65) libraries.

### Strains of *S. elongatus*, growth conditions and ciprofloxacin titration

Wild-type strain of *S. elongatus* PCC 7942 was obtained from Invitrogen. Strains with deletion and overexpression of *ago* gene were obtained previously using standard protocols (25). Cyanobacterial strains were maintained in liquid BG11 with shaking or on solid BG11 plates (66) under continuous light conditions (∼250 μE m^−2^ s^−1^) at 30°C with appropriate antibiotics (10 μg/ml spectinomycin) if needed.

Dense cultures of *S. elongatus* wild-type and mutant strains were inoculated into 800 μl of fresh BG11 medium supplemented with 0-35 ng/ml ciprofloxacin and spectinomycin (in the case of the mutant strains) in 48-well plates (Eppendorf). The plates were incubated at 300 rpm at 30°C under continuous light conditions and cell density was monitored by measuring OD_750_ every 12 hours. One biological replicate was performed for titration and three independent biological replicates were performed for growth experiment with 0 and 10 ng/ml ciprofloxacin. Cells were harvested for microscopy 48 hours after inoculation.

### Cell microscopy

*E. coli* cells were visualized using acridine orange staining. Sterilized slide was fixed with 95% ethanol for 2 minutes. Excess ethanol was drained, and slide is allowed to air-dry. The bacterial culture was placed onto the slide, dried and fixed. Slide was flooded with acridine orange stain for 2 minutes, rinsed thoroughly with tap water, and allowed to air-dry. Cell pictures were taken on a ZEISS LSM 900 confocal laser scanning microscope (Carl Zeiss) with a 63^×^ oil immersion objective and with 100-μm confocal pinhole aperture. Cell pictures of *S. elongatus* were taken similarly but using chlorophyll autofluorescence, which was excited at 488 nm and recorded using a 650 nm long-pass filter. The obtained pictures were processed using the ZEN Microscopy software (Carl Zeiss).

### Gyrase knockdown with dCas9

To knockdown *gyrA* gene, a plasmid carrying the pA15 origin, chloramphenicol resistance and dCas9 genes, and sgRNA was assembled. The 20 nt guide sequence in sgRNA corresponding to the *gyrA* gene (5’-AGCTCTTCCTCAATGTTGAC-3’) was chosen based on the algorithm developed in (67). *E. coli* Bl21(DE3) strains containing pBAD plasmids encoding or lacking pAgos were transformed with the dCas9 plasmid and grown under standard conditions. The drop of the *gyrA* expression level was measured by quantitative PCR using primers corresponding to the *gyrA* gene (qPCR_gyrA_for 5’-TTATGACACGATCGTCCGTATG and qPCR_gyrA_rev 5’-TTCCGTGCCGTCATAGTTATC) and the *rpoB* gene for normalization (rpoB_Fw1 5’-ATGGTTTACTCCTATACCGAGAAAAAAC and rpoB_Rv1 5’-TATTGCAGCTCGGAATTACCG).

## AUTHOR CONTRIBUTIONS

Andrey Kulbachinskiy and Alexei Aravin conceived the study, Anna Olina performed experiments, Aleksei Agapov performed bioinformatic analysis of small DNA libraries, Denis Yudin made the original discovery of the chromosomal specificity of pAgos under supervision of Anton Kuzmenko and performed initial analysis of small DNAs, Anna Olina and Aleksei Agapov prepared the figures, Andrey Kulbachinskiy wrote the manuscript with contributions from all the authors.

## ACKNOWLEDGMENTS

We thank Daria Esyunina for continued advice on this study, Phillip Zamore, Samson M. Jolly and Dmitry Sutormin for helpful discussions. This work was supported by grant 19-14-00359 of the Russian Science Foundation to Daria Esyunina. The authors declare no competing interests.

## DATA AVAILABILITY

The smDNA sequencing datasets generated in this study are available from the Sequence Read Archive (SRA) database under bioproject number PRJNA878808. The code used for data analysis is available at the GitHub repository at https://github.com/AlekseiAgapov/SeAgo_LrAgo. All primary data are available from the corresponding author upon request.

