## Supplemental Figures S1-S4 and Table S1 for "Bacterial Argonaute proteins aid cell division in the presence of topoisomerase inhibitors in *Escherichia coli*"

### SUPPLEMENTARY INFORMATION

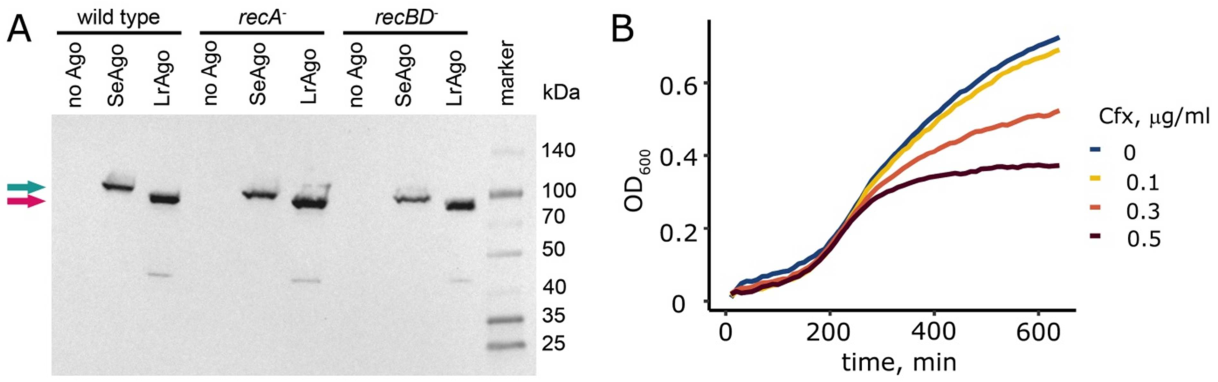


**Fig. S1.** (A) Analysis of pAgo expression in *E. coli* by Western blotting. Protein samples were obtained from the wild-type or *rec*-minus *E. coli* strains at 4.5 hours of growth. The proteins were separated by 4-20% gradient denaturing SDS-PAGE, transferred to a nitrocellulose membrane and visualized by Western-blotting with His-tag specific antibodies (see Materials and Methods for details). The green arrow points to SeAgo, the pink one indicates LrAgo. (B) Analysis of *E. coli* growth at sublethal concentrations of ciprofloxacin. Wild-type *E. coli* were grown at 30 °C in a plate reader with indicated concentrations of ciprofloxacin (0, 0.1, 0.3 or 0.5 μg/ml). Averages from 3 biological replicates are shown.

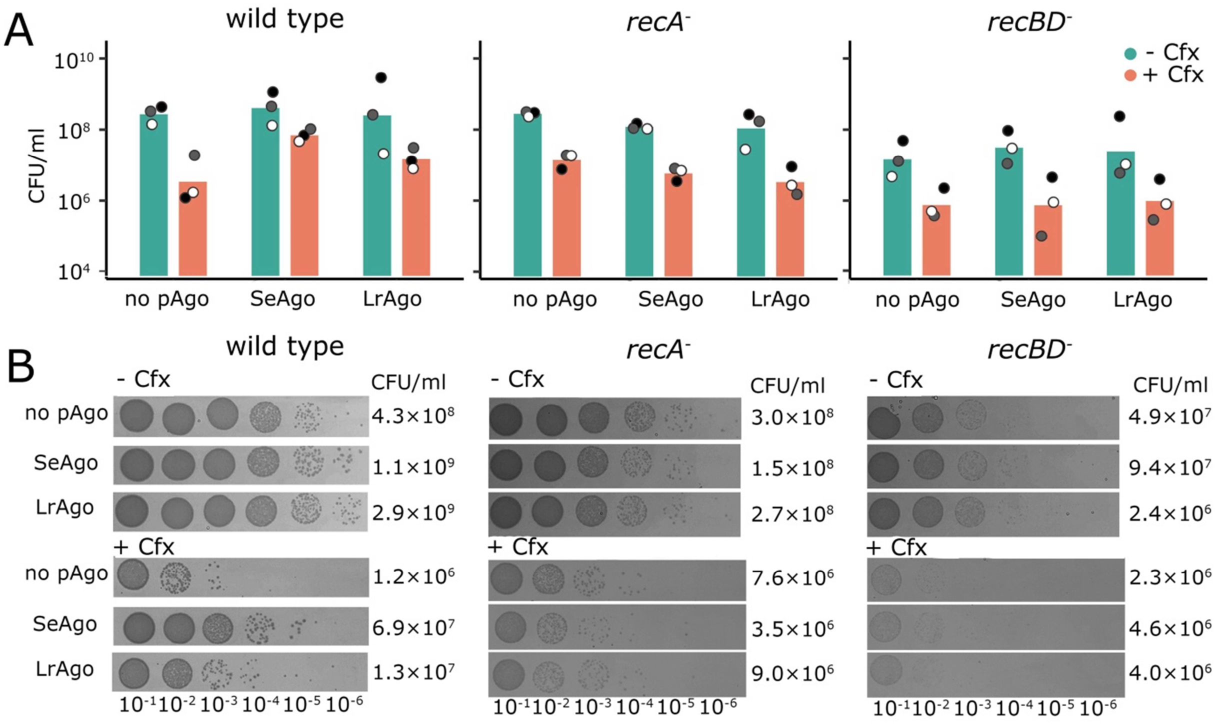


**Fig. S2. Analysis of the number of viable cells in *E. coli* strains lacking or containing pAgos in the absence and in the presence of ciprofloxacin.** (A) The numbers of colony forming units (CFU) in indicated *E. coli* strains. The samples were taken from *E. coli* cultures grown for 4.5 hours in the absence or in the presence of ciprofloxacin (Fig. 2), and CFU numbers were determined by plating their serial dilutions on LB agar plates without ciprofloxacin. Averages from three independent biological replicates are shown; individual data points from each measurement are indicated. (B) Representative LB plates for *E. coli* strains grown in the absence (top) or in the presence (bottom) of ciprofloxacin. The CFU numbers calculated for each strain are shown on the right.

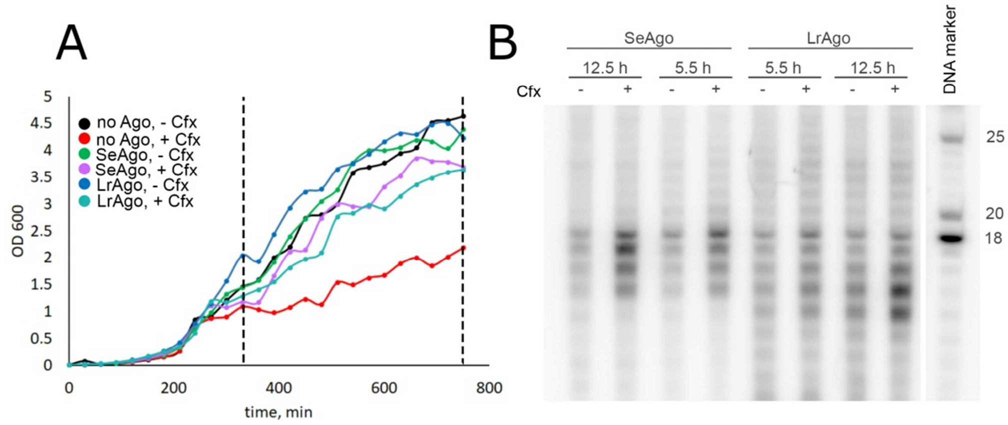


**Fig. S3. Analysis of smDNA libraries from *E. coli* strains expressing pAgos.** (A) Preparative-scale cultivation of *E. coli* strains lacking of containing pAgos (SeAgo, top; LrAgo, bottom). The cultures were grown in 0.5 liter of LB in the absence and in the presence of ciprofloxacin (0.3 μg/ml) and $OD_{600}$ was monitored each 30 minutes. The dashed lines indicate 5.5 h and 12.5 h time points used for purification of pAgo-associated smDNAs. (B) Analysis of smDNAs purified from pAgos. SmDNAs isolated from pAgos after one-step purification (using $Co^{2+}$-affinity resin) were treated with alkaline phosphatase to remove pre-existing 5'-phosphates, labeled with γ-$P^{32}$-ATP and polynucleotide kinase and separated by 19% denaturing urea PAGE. The marker lane (M) contains 5'-labeled DNA oligonucleotides of indicated lengths.

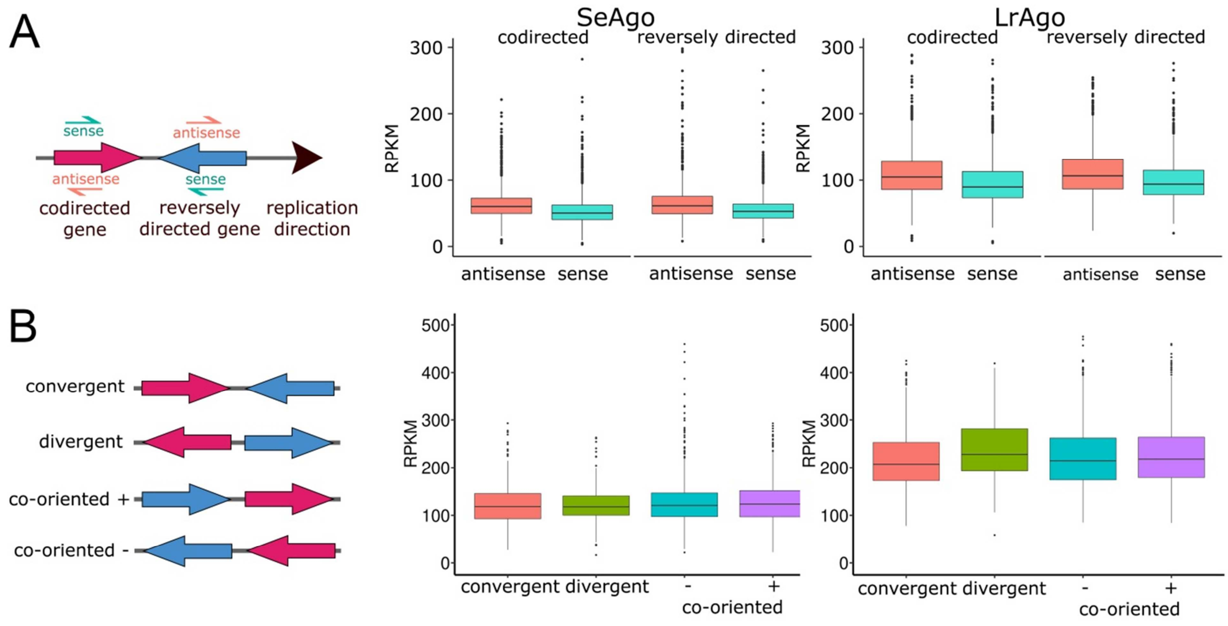


**Fig. S4. Analysis of smDNA distribution relative to transcription units in *E. coli*.** (A) Densities of smDNAs within genes co-directed or reversely directed relative to replication. For each gene, the amounts of smDNAs were calculated independently for the sense and antisense DNA strands, normalized by the gene length and expressed as RPKM (reads per kilobase per million reads in the library). The data were independently averaged for each of the four types of orientation (sense and antisense strands for co-directed and reversely directed genes. (B) Densities of smDNAs in intergenic regions for convergent, divergent, and co-oriented gene pairs (located either in the plus or in the minus genomic strands). Only <500 bp intergenic regions were taken into account. SmDNA numbers were independently averaged for each of the four types of gene orientation (convergent, divergent, and two directions of co-oriented genes), normalized by the length of each intergenic region and expressed in RPKM. The data are shown as box plots (center line, median; box limits, upper and lower quartiles; whiskers, 1.5x interquartile range).

**Table S1. Small DNA libraries obtained and analyzed in this study.** The data are available from the SRA database, bioproject number PRJNA878808

| library number | library name | pAgo | ciprofloxacin | time, h | SRA accession | reads mapped to genome | reads mapped to genome, % | reads mapped to plasmid | reads mapped to plasmid, % | enrichment in plasmid coverage* |
|---|---|---|---|---|---|---|---|---|---|---|
| 1 | Se_5.5_nocip_rep_1 | SeAgo | - | 5.5 | SRR21504308 | 368873 | 81.6 | 83301 | 18.4 | 11.4 |
| 2 | Se_5.5_nocip_rep_2 | SeAgo | - | 5.5 | SRR21508618 | 488441 | 79.3 | 127566 | 20.7 | 12.8 |
| 3 | Se_5.5_cip_rep_1 | SeAgo | 0.3 μg/ml | 5.5 | SRR21508717 | 1509243 | 88.4 | 197824 | 11.6 | 7.2 |
| 4 | Se_5.5_cip_rep_2 | SeAgo | 0.3 μg/ml | 5.5 | SRR21508866 | 1038304 | 87.2 | 152951 | 12.8 | 8 |
| 5 | Se_12.5_nocip_rep_1 | SeAgo | - | 12.5 | SRR21508879 | 791799 | 78.9 | 211831 | 21.1 | 13.1 |
| 6 | Se_12.5_nocip_rep_2 | SeAgo | - | 12.5 | SRR21508723 | 675406 | 77.7 | 193981 | 22.3 | 13.8 |
| 7 | Se_12.5_cip_rep_1 | SeAgo | 0.3 μg/ml | 12.5 | SRR21508709 | 2152542 | 87.1 | 318401 | 12.9 | 8 |
| 8 | Se_12.5_cip_rep_2 | SeAgo | 0.3 μg/ml | 12.5 | SRR21508927 | 2920406 | 86.4 | 460114 | 13.6 | 8.4 |
| 9 | Lr_5.5_nocip_rep_1 | LrAgo | - | 5.5 | SRR21508877 | 1266921 | 94.3 | 75910 | 5.7 | 3.5 |
| 10 | Lr_5.5_nocip_rep_2 | LrAgo | - | 5.5 | SRR21508707 | 1984106 | 94.8 | 108631 | 5.2 | 3.2 |
| 11 | Lr_5.5_cip_rep_1 | LrAgo | 0.3 μg/ml | 5.5 | SRR21508916 | 1290785 | 92.5 | 105237 | 7.5 | 4.7 |
| 12 | Lr_5.5_cip_rep_2 | LrAgo | 0.3 μg/ml | 5.5 | SRR21508919 | 1265045 | 91.3 | 121100 | 8.7 | 5.4 |
| 13 | Lr_12.5_nocip_rep_1 | LrAgo | - | 12.5 | SRR21508925 | 1196750 | 86.0 | 194585 | 14.0 | 8.6 |
| 14 | Lr_12.5_nocip_rep_2 | LrAgo | - | 12.5 | SRR21508926 | 1003480 | 87.4 | 145080 | 12.6 | 7.8 |
| 15 | Lr_12.5_cip_rep_1 | LrAgo | 0.3 μg/ml | 12.5 | SRR21508713 | 2974923 | 83.1 | 606222 | 16.9 | 10.4 |
| 16 | Lr_12.5_cip_rep_2 | LrAgo | 0.3 μg/ml | 12.5 | SRR21508921 | 1556966 | 84.5 | 285548 | 15.5 | 9.6 |

* The enrichment of plasmid-derived smDNAs is calculated as a ratio of the observed number of plasmid-mapped reads to the expected number of plasmid mapped reads. The expected number of reads = (genome-mapped reads + plasmid-mapped reads)×(plasmid size × copy number)/( genome size + plasmid size × copy number)
